# The effect of clinically relevant beta-lactam, aminoglycoside, and quinolone antibiotics on bacterial extracellular vesicle release from *E. coli*

**DOI:** 10.1101/2023.11.22.568081

**Authors:** Panteha Torabian, Navraj Singh, James Crawford, Gabriela Gonzalez, Nicholas Burgado, Martina Videva, Aidan Miller, Janai Perdue, Milena Dinu, Anthony Pietropaoli, Thomas Gaborski, Lea Vacca Michel

## Abstract

Sepsis, a leading cause of death in hospitals, can be defined as a dysregulated host inflammatory response to infection, which can lead to tissue damage, organ failure, and cardiovascular complications. Although there is no cure for sepsis, the condition is typically managed with broad spectrum antibiotics to eliminate any potential bacterial source of infection. However, a potential side-effect of antibiotic treatment is the enhanced release of bacterial extracellular vesicles (BEVs). BEVs are membrane-bound nanoparticles produced by a variety of mechanisms, one of which includes the pinching-off of the outer membrane (in Gram-negative bacteria) to enclose proteins and other biological molecules for transport and intercellular communication. Some of the Gram-negative EV cargo, including Peptidoglycan associated lipoprotein (Pal) and Outer membrane protein A (OmpA), have been shown to induce both acute and chronic inflammation in host tissue. We hypothesize that antibiotic concentration and its mechanism of action can have an effect on the amount of released BEVs, which could potentially exacerbate the host inflammatory response. In this study, we evaluated nine clinically relevant antibiotics for their effect on EV release from *Escherichia coli*. EVs were characterized using immunoblotting, nanoparticle tracking analysis, and transmission electron microscopy. Several beta-lactam antibiotics caused significantly more EV release, while quinolone and aminoglycosides caused relatively less vesiculation. Further study is warranted to corroborate the correlation between an antibiotic’s mechanism of action and its effect on EV release, but these results underline the importance of antibiotic choice when treating sepsis patients.

## Introduction

According to the International Society for Extracellular Vesicles (ISEV), extracellular vesicles (EVs) are membrane-enclosed nanobodies that are naturally released from all cell types and lack a functional nucleus [1]. Bacterial extracellular vesicles (BEVs) are generated by both Gram-positive (GP) and Gram-negative (GN) bacterial cells, containing a multitude of cellular components, including intracellular soluble and membrane-associated proteins and nucleic acids. These components originate from the parent bacterium from which they derive, and their inclusion in BEVs are largely dependent upon the mechanism of BEV biogenesis. There are two major types of BEVs generated by GN bacteria: outer membrane vesicles (OMVs) that pinch off from the outer membrane and apoptotic bodies (ApoBDs) that form during the final stages of apoptosis. OMVs are generally smaller in size (20–250 nm in diameter) compared to ApoBDs (500-5,000nm in diameter) [1], [2].

Understanding how and why antibiotics enhance the release of bacterial EVs is crucial, especially in the context of sepsis patients who often receive broad-spectrum antibiotics as a first line of treatment. OMVs containing LPS, virulence factors, adhesins, and lipoproteins have been implicated in initiating inflammation during the transition from infection to sepsis [3]–[5]. They also play a complex role in endothelial activation and can induce cardiac injury, worsening patient outcomes in sepsis [3]–[5].

The immunomodulatory molecules contained within OMVs, including LPS and other outer membrane proteins, are thought to interact with host cells through several different mechanisms, such as activating host immune cells via TLRs (e.g., TLR4), triggering the release of pro- and anti-inflammatory cytokines, and delivering bacterial content into host cells [6]–[8] Additionally, due to their small size, OMVs are capable of self-entry deep into host tissues, engaging both the innate and adaptive immune systems and resulting in longer-term, chronic responses and inflammatory pathologies [5], [9]–[11].

Here, we test the hypothesis that certain classes of antibiotics enhance BEV release from GN bacteria, specifically *Escherichia coli* (*E. coli*), more so than other antibiotics. We developed this hypothesis, in part, based on our own studies, as well as other studies that have looked at the effect of antibiotics on GN BEV release [12] [13]. For example, scientists have shown that treatment of *Shigella dysenteriae* with mitomycin increases BEV production, while fosfomycin, ciprofloxacin, and norfloxacin did not have a significant effect on BEV release [13]. Another study showed that gentamicin treatment of *Pseudomonas aeruginosa* resulted in a threefold increase in BEV production [14]. A third study demonstrated that ciprofloxacin, meropenem, fosfomycin, and polymyxin B increased the production of BEVs in two strains of *E. coli* (O104:H4 and O157:H7) [14].

In this study, we tested nine clinically relevant antibiotics for their effect on EV release from a clinical strain of *E. coli* (*E. coli* K1 RS218). These antibiotics are commonly used to treat sepsis patients at the University of Rochester Medical Center (URMC, Rochester, NY): six beta-lactam antibiotics (ceftriaxone, piperacillin, imipenem, ertapenem, meropenem, cefepime), two aminoglycoside antibiotics (tobramycin and amikacin), and one quinolone antibiotic (ciprofloxacin). A brief summary of each antibiotic’s mechanism of action is described in Table 1. We performed our experiments using two dosage strategies. In the first set of experiments, we employed the antibiotics at twice their minimum inhibitory concentration (2MIC), determined using the broth dilution method. In the second set of experiments, we used the antibiotics at concentrations proportional to those commonly used in the clinic. Bacterial EVs were quantified and characterized using immunoblotting (western blot), nanoparticle tracking analysis (NTA), and transmission electron microscopy (TEM). We observed that the beta-lactam antibiotics resulted in greater amounts of BEV release compared to aminoglycoside and quinolone antibiotics. We also determined that there is variability in the amount of BEVs released among the different beta-lactam antibiotics, perhaps due to subtle differences in their specific inhibitory mechanisms of action.

**Table 1.** Mechanism of action of clinically relevant antibiotics, six beta-lactam antibiotics (ceftriaxone, piperacillin, imipenem, ertapenem, meropenem, cefepime), two aminoglycoside antibiotics (tobramycin and amikacin), and one quinolone antibiotic (ciprofloxacin). Generally, beta-lactam antibiotics target cell wall synthesis, aminoglycosides target protein synthesis, and quinolones target DNA synthesis.

| Antibiotic | Mechanism of Action |
| --- | --- |
| **Ceftriaxone** | Inhibition of mucopeptide synthesis in the bacterial cell wall [15] |
| **Piperacillin** | Inhibition of penicillin-binding proteins, leading to disruption of bacterial cell wall cross-linkages [16] |
| **Imipenem** | Inhibition of bacterial cell wall synthesis by binding to and inactivating relevant transpeptidases, known as penicillin binding proteins (PBPs), with specific high affinity to PBPs-2, -4, and -5 in *E. coli* [17] |
| **Ertapenem** | Inhibition of the elongation and reinforcement of the peptidoglycan component of the bacterial cell wall by binding to penicillin-binding proteins (PBPs) [18] |
| **Meropenem** | Inhibition of cell wall synthesis by penetrating the cell wall and binding to penicillin-binding-protein (PBP), with specific high affinity to PBPs 2 in *E. coli* [19] |
| **Cefepime** | Inhibition of cell wall synthesis, leading to lysis, by binding to PBPs; cefepime's zwitterionic nature aids in the rapid penetration of the GN outer membrane [20] |
| **Tobramycin** | Inhibition of protein synthesis by binding to the bacterial 30S and 50S ribosome, preventing formation of the 70S complex; as a result, mRNA cannot be translated into protein [21] |
| **Amikacin** | Inhibition of protein synthesis by binding to bacterial 30S ribosomal subunits, interfering with mRNA binding and tRNA acceptor sites [22] |
| **Ciprofloxacin** | Inhibition of DNA replication by inhibiting bacterial DNA topoisomerase and DNA-gyrase [23], [24] |

## Results

### Antibiotic concentrations

We used two different strategies to determine the antibiotic concentrations for our experiments. For the first set of experiments, we determined the minimum inhibitory concentration (MIC) for each antibiotic (Table 2), which is the lowest concentration of antibiotic that inhibited visible growth of bacteria (*E. coli* strain K1 RS218). For the first set of experiments, we considered the general intravenous dosages for each antibiotic commonly used by physicians at URMC. The “clinical concentrations” were determined for each antibiotic to be roughly proportional to the clinical dosage, as described in the methods section and in Table 2. In the second set of experiments, we incubated the bacteria with twice the MIC (2MIC) of each antibiotic for 3.5 hours prior to isolating EVs using ultracentrifugation.

**Table 2.**
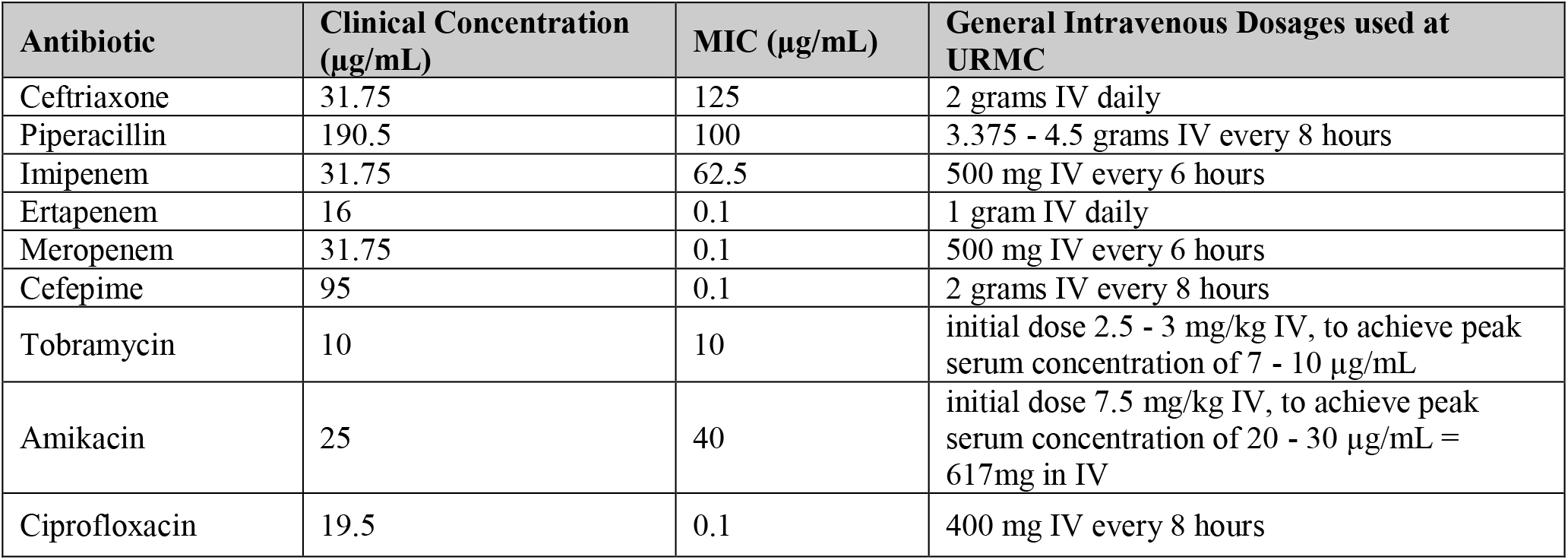
Clinical concentrations and Minimum Inhibitory Concentrations (MIC) of each antibiotic. Clinical concentrations (μg/mL) were calculated to be proportional to general intravenous dosages commonly used at URMC (Rochester, NY), and MICs were determined using the broth dilution method (μg/mL). For experiments, twice the MICs (2MIC) of antibiotics were used.

### Protein Quantification

EVs were isolated from *E. coli* cells, post-incubation with either clinical concentrations or 2MIC of antibiotics (or no antibiotic, as a control). The isolated EVs were quantified using immunoblotting with antibodies to *E. coli* peptidoglycan associated lipoprotein (Pal) and outer membrane protein A (OmpA). In some cases, particularly when no antibiotic was used, only faint bands were detected, in which case we assumed that although EVs were likely produced, the amount of isolated EVs was below our limit of detection. Pal and OmpA band volumes for both sets of experiments are shown in **Figure 1**. Overall, band volumes were greater for beta-lactam samples compared to aminoglycoside and quinolone samples. However, these differences are reduced in the 2MIC experiment.

**Figure 1.**
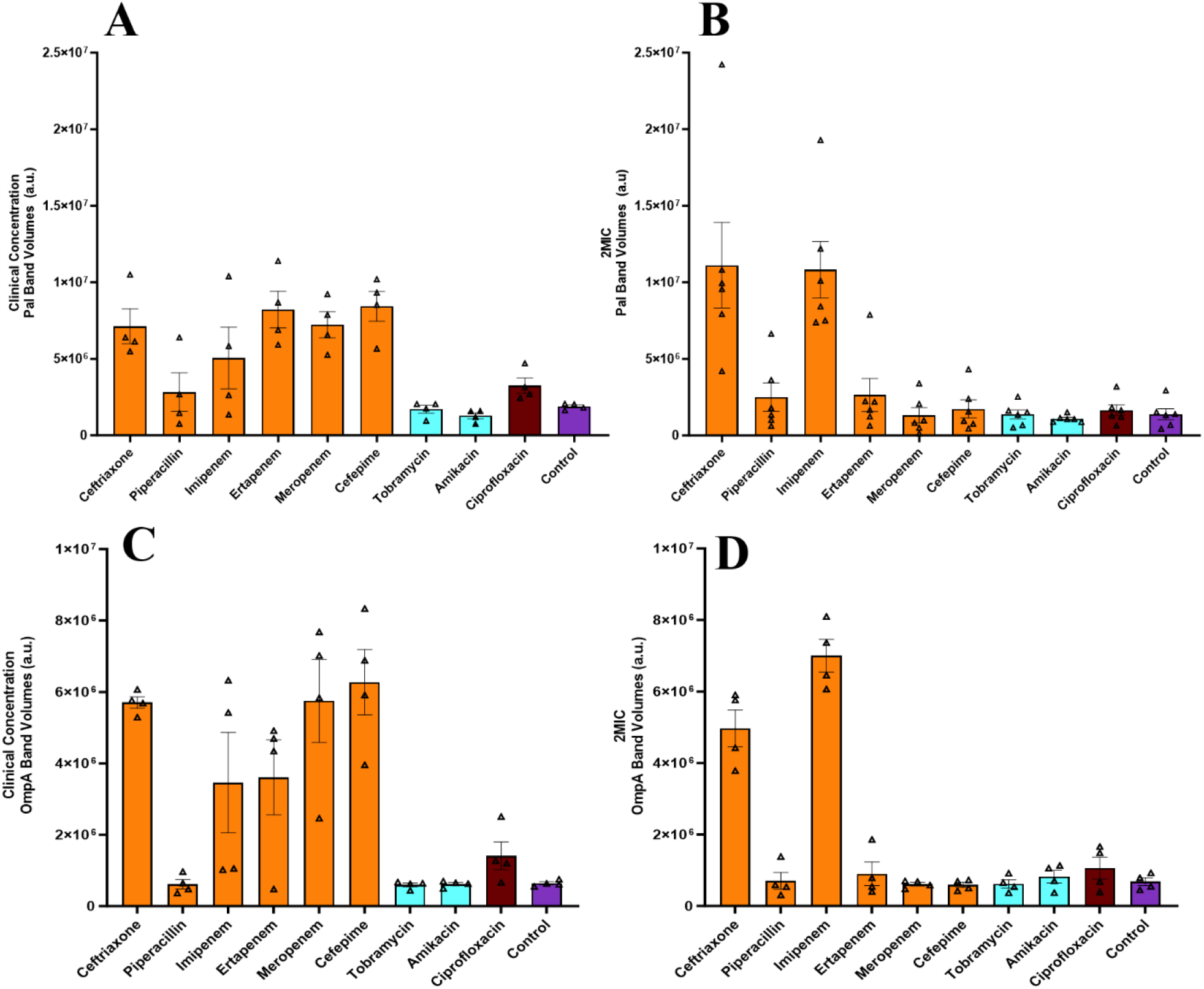
BEV estimates using *E. coli* antigen band volumes. The average Pal band volumes measured from immunoblots for clinical concentration experiments (A, n=4) and 2MIC experiments (B, n=6). The average OmpA band volumes measured from immunoblots for clinical concentration experiments (C, n=4) and 2MIC experiments (D, n=4). All individual data points are shown and represent independent biological replicates. Error bars represent the standard error of the mean (SEM), determined using GraphPad Prism.

### Nanoparticle tracking analysis

Nanoparticle Tracking Analysis (NTA) is a technique used to measure the size distribution and concentration (particle/mL) of nanoparticles in a liquid sample. NTA employs laser light scattering to individually track and analyze the Brownian motion of nanoparticles, estimating the size of the particles based on their diffusion properties. Results from the NTA experiments are shown in **Figure 2**. Overall, the BEV concentrations, as estimated by NTA, correlate well with the estimates determined using protein band volumes.

**Figure 2.**
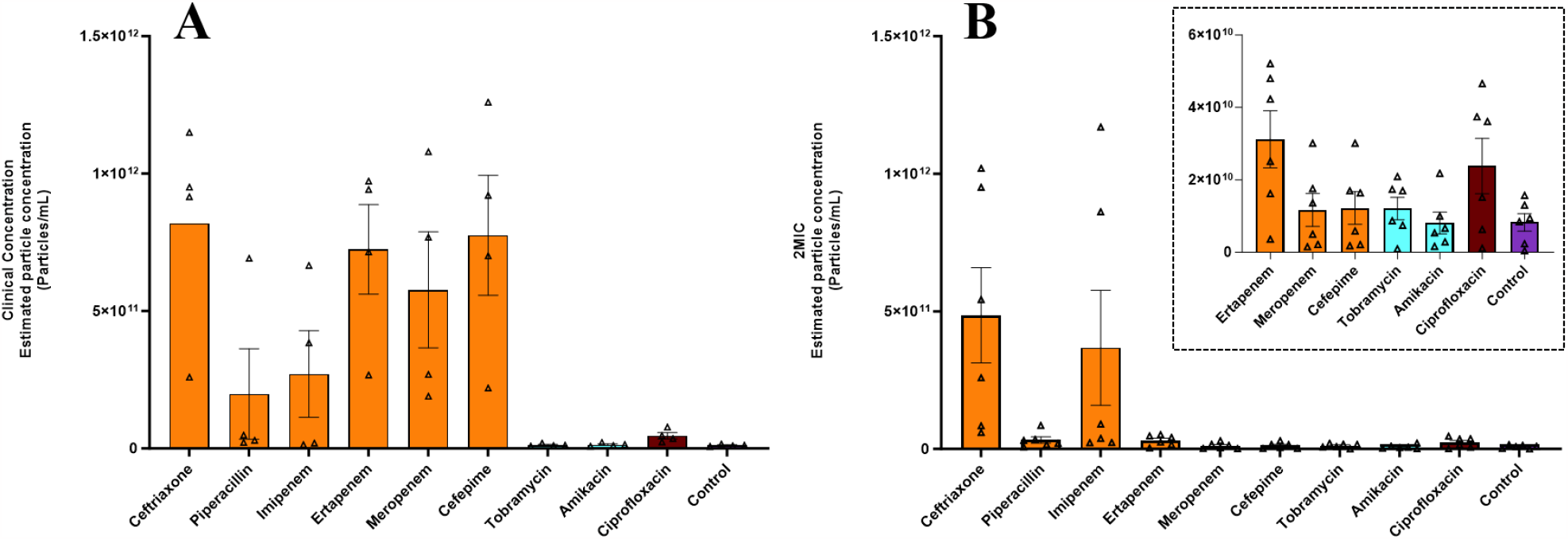
Average particle concentrations determined using nanoparticle tracking analysis (NTA). The average particle concentrations for clinical concentration experiments (A, n=4) and 2MIC experiments (B, n=6). All individual data points are shown and represent independent biological replicates. Error bars represent the standard error of the mean (SEM), determined using GraphPad Prism.

### Transmission Electron Microscopy

All BEV samples were prepared as described above for immunoblotting and NTA and then imaged with transmission electron microscopy (TEM). We selected a subset of antibiotics (ceftriaxone, cefepime, ciprofloxacin, amikacin) to test for their effect on BEV release. As expected, BEVs have a distinct bilayer membrane and sometimes exhibit a puckering effect, as is a common side effect from the TEM staining/drying process (**Figures 3 and 4**). We also determined the average particle size for each antibiotic, as shown in **Figure 5**.

**Figure 3.**
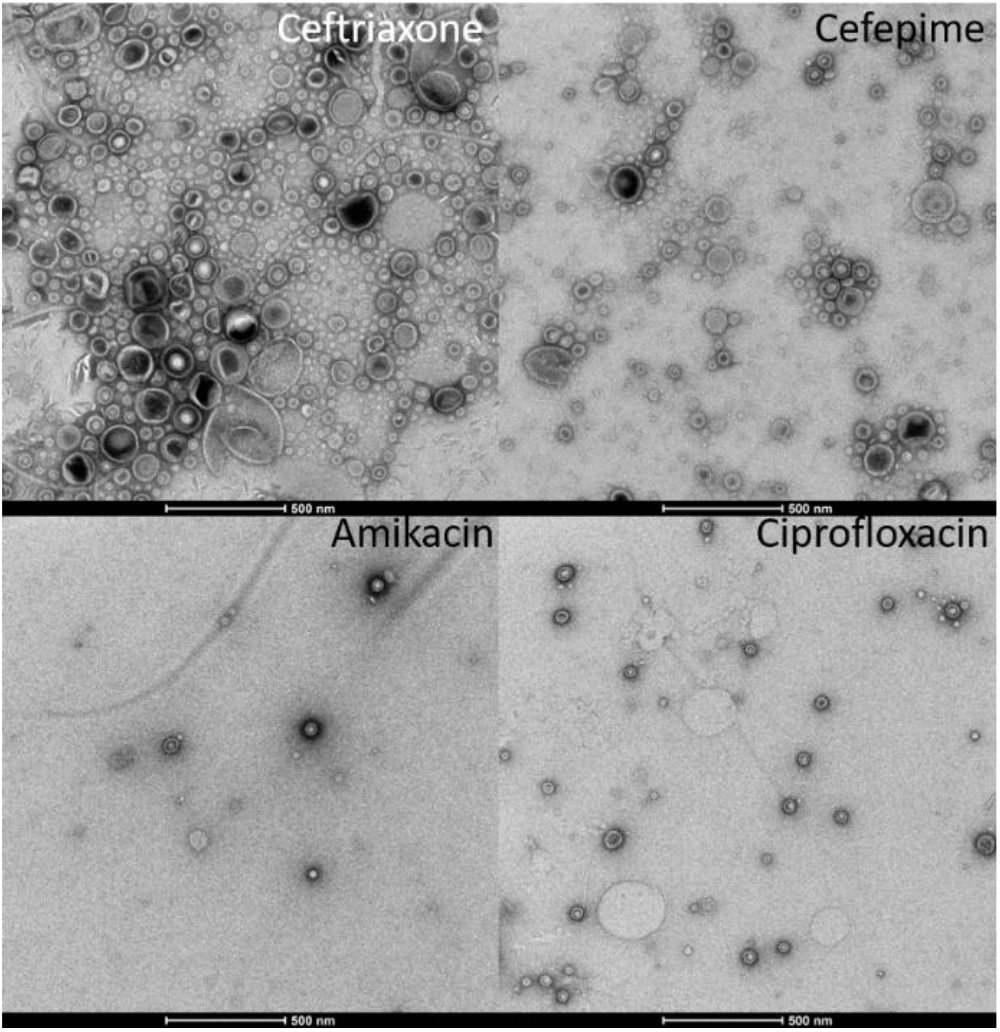
Transmission electron microscopy (TEM) images, clinical concentrations of antibiotics. TEM images of BEVs from *E. coli* incubated with clinical concentrations of antibiotics: ceftriaxone, cefepime, amikacin, and ciprofloxacin (scale bar: 500 nm).

**Figure 4.**
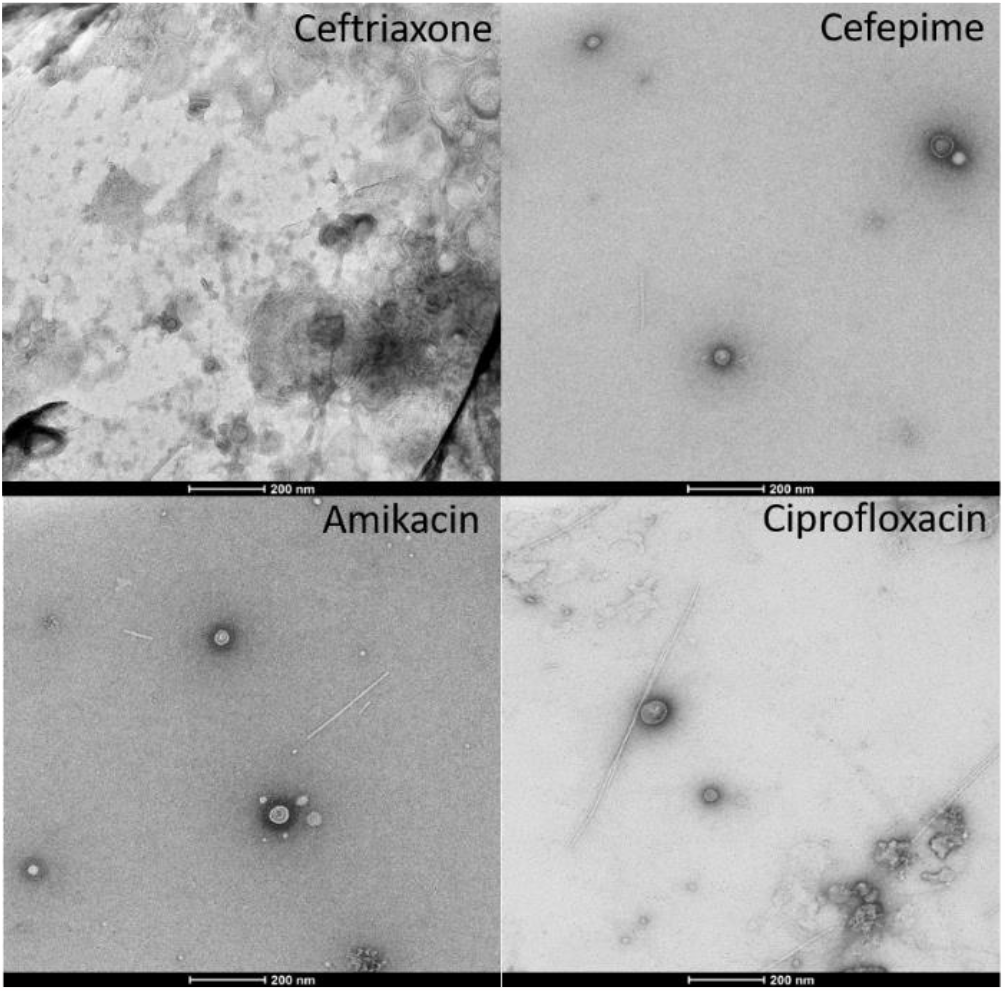
Transmission electron microscopy (TEM) images, 2MIC of antibiotics. TEM images of BEVs from *E. coli* incubated with 2MIC of antibiotics: ceftriaxone, cefepime, amikacin, and ciprofloxacin (scale bar: 200 nm).

**Figure 5.**
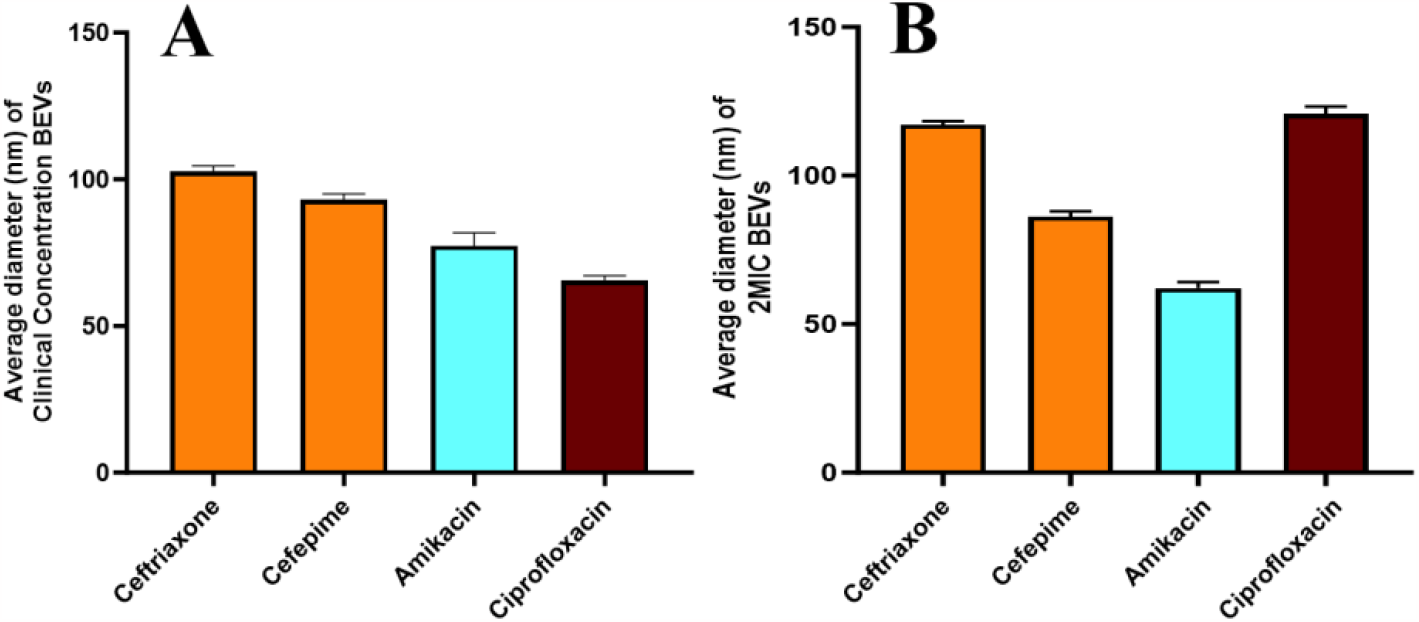
Transmission electron microscopy (TEM) image analysis. Average particle size (n=5) of BEVs, measured for each antibiotic using ImageJ software, released from *E. coli* incubated with clinical concentrations of antibiotics (A) and 2MIC of antibiotics (B). Error bars represent the standard error of the mean (SEM).

### Relative BEV release

From the results of the experiments described above, we determined the relative amounts of BEVs released from *E. coli* in the presence of antibiotics versus control (no antibiotics), according to Pal band volume measurements. As seen in **Figure 6**, beta-lactam antibiotics enhanced the release of BEVs from *E. coli* between 3 and 5-fold compared to control, while the one quinolone tested (ciprofloxacin) exhibited a small increase in BEV release compared to control. Aminoglycosides (tobramycin and amikacin) averaged similar BEV counts as seen for the control sample, suggesting that the aminoglycoside antibiotics do not enhance BEV production. Moreover, beta-lactam antibiotics induced about 2 to 4-fold more BEV release compared to the other groups, while the quinolone induced only 1 to 2-fold more BEVs compared to aminoglycosides (**Figure 7**). We performed a similar analysis, comparing BEVs for beta-lactam, aminoglycoside, and quinolone antibiotics compared to control (no antibiotics) and compared to each other, using NTA-estimated particle counts (**Supplementary Figures S1 and S2**). The ratios of BEVs between subgroups were much higher for NTA-estimated BEV counts, particularly for beta-lactam antibiotics, compared to those ratios determined by Pal band volumes; however, the trends were similar for both sets of data.

## Discussion

Bacterial extracellular vesicles (BEVs) have been implicated in both inflammatory and anti-inflammatory host responses and are thought to contribute to disease pathogenesis, due largely to their cargo, which can include toxic, virulent, and inflammatory molecules. Because of their inflammatory contributions, BEVs have been studied as potential therapeutic targets, vaccine delivery systems, and diagnostic biomarkers [12].

Here, we describe our investigation into the effect of clinically relevant antibiotics, at two biologically-significant concentrations, on BEV release from *E. coli*. We measured BEV release using two approaches: quantification of two protein antigens commonly found in *E. coli* BEVs, peptidoglycan-associated lipoprotein (Pal) and outer membrane protein A (OmpA), using immunoblotting and particle counts using nanoparticle tracking analysis (NTA). We also characterized the BEVs using transmission electron microscopy (TEM). Although there was some variability in the data, the results from these approaches converged on several common themes.

First, when *E. coli* bacteria are incubated with antibiotics at 2MIC or clinical concentrations, beta-lactam antibiotics induce approximately 4-fold the amount of BEV release compared to the aminoglycosides, 2 to 3-fold compared to the quinolone antibiotic, and 3 to 5-fold compared to no antibiotics (**Figures 6 and 7**). The aminoglycosides released similar amounts of BEVs to control, suggesting that the two aminoglycoside antibiotics used in this study do not enhance BEV production or release. The relative numbers described here were calculated from average Pal band volumes, which results in several caveats. For example, Pal band volumes will only reflect BEV counts if their expression levels are consistent per BEV across conditions. However, doing a similar analysis using NTA particle counts yielded similar trends, suggesting that Pal expression was consistent across BEVs. A second caveat to consider is the limitation of testing only one quinolone antibiotic and two aminoglycoside antibiotics compared to six beta-lactam antibiotics. As one can see from the beta-lactam data, BEV release varies between the individual beta-lactam antibiotics, perhaps due to their unique mechanisms of action. If we tested several more quinolones or aminoglycosides, we could possibly get significantly varied results.

**Figure 6.**
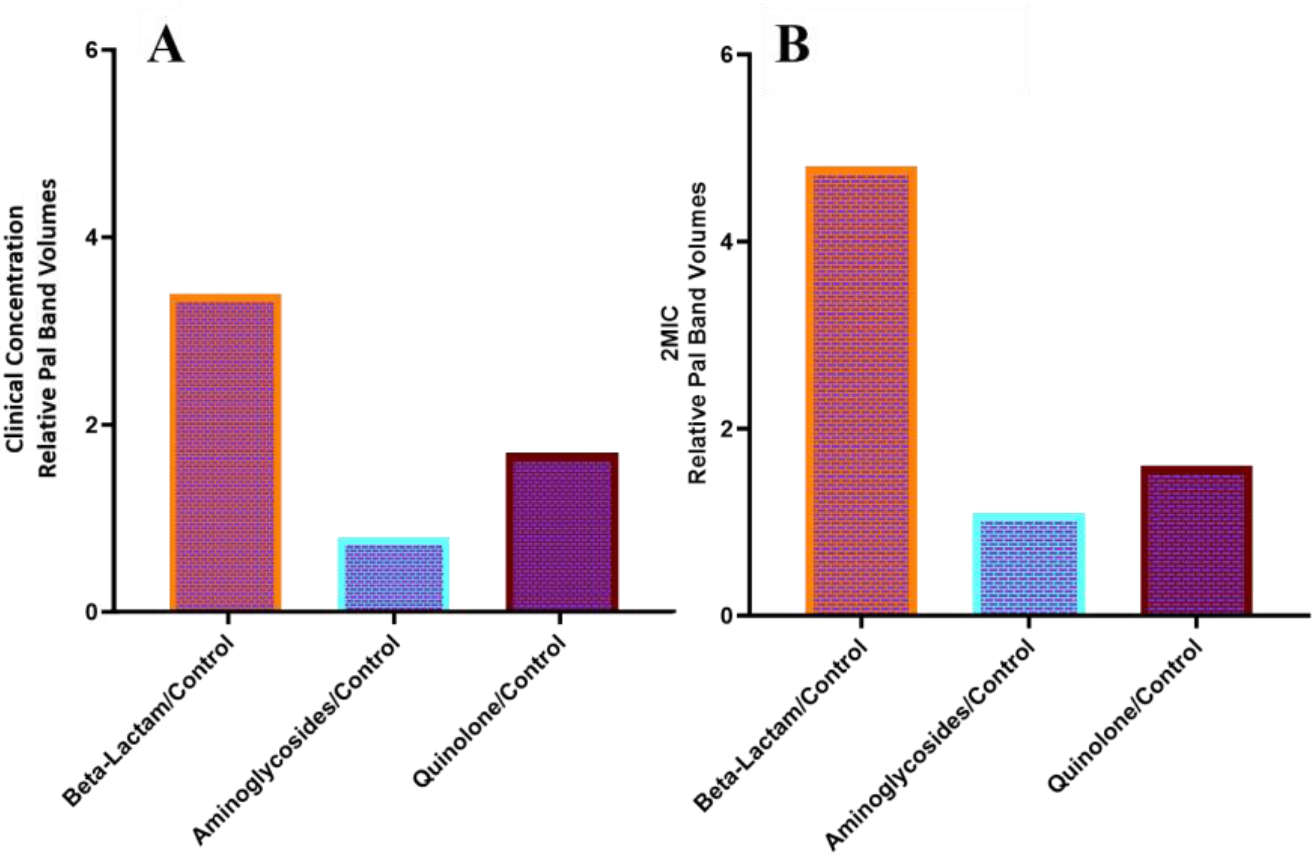
Relative BEV counts determined from average Pal band volumes. Relative BEV counts were determined using the average Pal band volumes for each class of antibiotic compared to control (no antibiotic) for clinical concentrations (A) and 2MIC (B).

**Figure 7.**
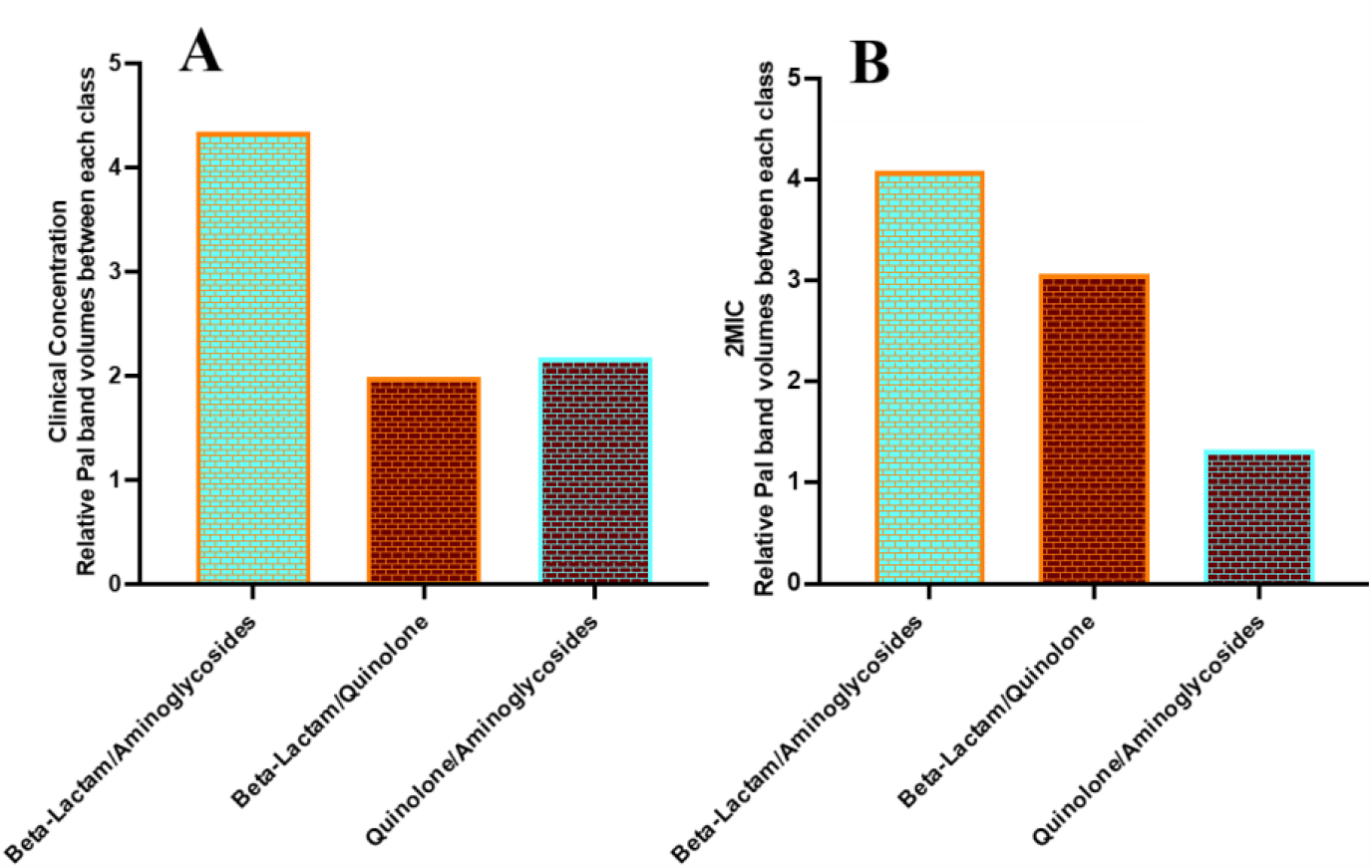
Relative BEV counts between antibiotic classes. Relative BEV counts, as determined by average Pal band volumes, for beta-lactam antibiotics vs aminoglycosides, beta-lactam antibiotics vs quinolone, and quinolone vs aminoglycosides at clinical concentrations (A) and 2MIC (B).

Third, incubating *E. coli* with antibiotics at twice their MICs reduces the variability between antibiotic types, with the exception of ceftriaxone and imipenem, as measured by immunoblotting and NTA (**Figures 1 and 2**). However, the two aminoglycosides, at both 2MIC and clinical concentrations, consistently released BEVs at similar levels to those of the negative control (**Figures 1 and 2**), where no antibiotics were present, again suggesting that the aminoglycoside antibiotics do not enhance BEV production.

A fourth common theme was that ceftriaxone was always among the greatest BEV-producing antibiotics. Minimum inhibitory concentrations of antibiotics were determined by incubating *E. coli* bacteria in LB media with serial dilutions of each antibiotic overnight; the lowest concentration of antibiotic that prevented bacterial growth was designated as that antibiotic’s MIC. However, it has been reported that ceftriaxone may degrade quickly when in solution (within four hours), perhaps resulting in our overestimation of ceftriaxone’s MIC [25]. The TEM images, which show an excess of bacterial debris (more so than for the other antibiotics), corroborate the hypothesis that our MIC for ceftriaxone is an overestimate. Alternate studies have determined MICs for ceftriaxone with *E. coli* to be closer to 8μg/mL,15-fold lower than our estimated 125 μg/mL [26]. As seen in the clinical concentration data, when incubated with lower doses of ceftriaxone (31.75 μg/mL), *E. coli* produces BEVs at levels comparable with the other beta-lactam antibiotics. We also noticed that piperacillin produced lower BEV levels compared to the other beta-lactam antibiotics. Piperacillin is often used in the clinic in combination with tazobactam, a beta-lactamase inhibitor, suggesting that the activity of piperacillin alone (as used in this study) was likely inhibited by beta-lactamases excreted by the *E. coli* bacteria [27].

The last common theme that we drew from this study was that administering antibiotic doses beyond twice their minimum inhibitory concentrations (2MIC) leads to extensive BEV release. Specifically, we observed much greater BEV production with clinical concentrations of ertapenem, meropenem, and cefepime compared to 2MIC of the same antibiotics, where clinical concentrations were estimated to be >100-fold higher than the 2MICs (**Figures 1 and 2**). These results align with emerging evidence of antibiotics disrupting bacterial equilibrium and inducing stress-driven BEV production [28].

In most sepsis cases where antibiotics are the first line of treatment, the results of this study emphasize the intricate and complex nature of antibiotic dosing, considering the potential impact of BEVs and their ability to contribute to inflammation and pathogenicity. However, we acknowledge that most sepsis patients are prescribed broad-spectrum antibiotics, because the specific bacterial cause of infection is unknown, further complicating matters and requiring high doses of antibiotics in order to eliminate a broad range of possible bacterial pathogens. However, our study highlights a potential unfavorable outcome of this treatment strategy, which is the enhancement of BEV production, which can contribute to or exacerbate sepsis-related inflammation or, due to BEV’s ability to enter deep into human tissue, result in longer-term chronic responses to infection and inflammatory pathologies.

## Experimental procedures

### Bacterial Strain

*E. coli* strain K1 RS218 was a gift from Dr. Kwang Sik Kim (Johns Hopkins Medical Center). *E. coli* were cultured in LB broth, shaking between 120-200rpm and incubated at 37°C.

### Determination of minimum inhibitory concentrations

Six beta-lactam antibiotics, two aminoglycosides, and one quinolone antibiotic were selected based on their clinical significance, accessibility, and usage to treat sepsis patients at URMC. The minimum inhibitory concentration (MIC) of each antimicrobial agent was determined using the broth-dilution assay. In summary, *E. coli* was cultured in LB overnight. The growth was used to inoculate individual cultures (in test tubes) with serially diluted antibiotics in LB. A negative control culture contained no antibiotic. The cultures were incubated overnight at 37°C, shaking at 180rpm, and then checked visually for the lowest concentration of antibiotic with no visible cloudiness.

### Clinical concentration calculation

To estimate the “clinical concentrations” of antibiotics, we used an average weight of 70 kg per person. In the clinic, a loading dose of 3mg/kg of tobramycin is given initially, followed by maintenance dosing to reach a peak serum concentration of 7-10 μg/ml. Therefore, we estimated 210 mg of Tobramycin was given to the patient every 8 hours (3mg/kg * 70kg = 210mg), or 630mg given to the patient per day. We assumed 630mg of tobramycin was administered per day to get a peak serum concentration of 10 μg/ml. Therefore, all clinical concentrations were calculated using the following formula: X grams administered per day divided by 0.63 g x 10 μg/ml. We recognize that our “clinical concentrations” may be over-estimated, since they were based on a total tobramycin dose of 630 mg over 24 hours, which would be high for a 70 kg person. Nevertheless, these doses were normalized across all drugs tested and therefore apply equally to all antibiotic classes used. Therefore, our comparisons are valid and provide proof of principle that BEV release depends on antibiotic class.

| Antibiotics | Hospital Dosage | Grams/day | “Clinical Concentrations” as determined for these experiments |
| --- | --- | --- | --- |
| Ceftriaxone | 2 grams IV daily | 2 | 31.75 ug/ml |
| Piperacillin | 3.375 - 4.5 grams IV every 8 hours | 12 | 190.5 ug/ml |
| Imipenem | 500 mg IV every 6 hours | 2 | 31.75 ug/ml |
| Ertapenem | 1 gram IV daily | 1 | 16 ug/ml |
| Meropenem | 500 mg IV every 6 hours | 2 | 31.75 ug/ml |
| Cefepime | 2 grams IV every 8 hours | 6 | 95 ug/ml |
| Tobramycin | initial dose 3 mg/kg IV, to achieve peak serum concentration of 7 - 10 mcg/mL | 0.63 | 10 ug/ml |
| Amikacin | initial dose 7.5 mg/kg IV, to achieve peak serum concentration of 20 - 30 mcg/mL |  | 25 ug/ml |
| Ciprofloxacin | 400 mg IV every 8 hours | 1.2 | 19.1 ug/mL |
|  |  |  | *Use equation (X grams/0.63)*10 |

### EVs isolation and characterization

*E. coli* were cultured on LB agar at 37ºC overnight; colonies from the plate were used to inoculate a 50 mL growth, which was cultured overnight at 37ºC, shaking at 180 rpm. The small culture was used to inoculate two larger cultures (2x300mL), which were grown to log phase (optical density at 600nm ∼0.8). The two 300mL cultures were mixed and then aliquoted into ten sterilized 125mL flasks (50mL of culture in each flask). All antibiotic solutions were prepared, as described above at either 2MIC or their calculated clinical concentration, and then aliquoted into the corresponding flask. The *E. coli* cultures were incubated with antibiotic (or no antibiotic, for control) for 3.5 hours (37°C, 180 rpm). Cells were pelleted for 15 minutes at 5,000*xg*, and the supernatants were syringe filtered using 0.45μm filters and then concentrated from ∼50mL to ∼30 mL using 50kDa MWCO Amicon filters (Millipore). Extracellular vesicles were isolated by ultracentrifugation (one hour at 100,000*xg*, Beckman Coulter The Optima™ MAX-XP, TLA-120.1 Fixed-Angle Rotor), and resuspended in 200μl of phosphate buffered saline (PBS).

Samples were analyzed using standard SDS-PAGE (4-16% bis acrylamide, Precast gels, VWR) and immunoblotting techniques. For immunoblotting, we used rabbit polyclonal antibody anti-OmpA (∼38kDa; Antibody Research Corporation) or unpurified anti-Pal antisera from mice inoculated with purified recombinant non-lipidated Pal protein (∼21 kDa; contains a 6xHis-tag for purification; Rochester General Hospital). Immunoblots were developed using SuperSignal™ West Femto Maximum Sensitivity Substrate (ThermoFisher) and imaged using the BioRad ChemiDoc Imaging System. Band volumes were quantified using Biorad’s Quantity One software package.

### Statistical analysis and normalization

All experiments were performed four or six times (independent biological replicates), and the bar graphs are presented as mean values with error bars as standard error of the mean (SEM), calculated using GraphPad Prism 10.0 (GraphPad, San Diego, CA, USA). When the experiments were performed, all antibiotic samples were prepared at the same time (and from the same culture), but the BEV samples were run on two gels. Therefore, we normalized the band volumes from one gel to the other using the imipenem sample or purified Pal protein, where those samples were run on both gels and then normalized to be the same volume.

### TEM imaging

The BEVs were isolated from cultures grown with 2MIC of ceftriaxone, cefepime, amikacin, ciprofloxacin, or no antibiotic (control). 200 mesh copper grids coated with formvar/carbon film were glow discharged for 30s at 30 mA in a PELCO Easiglow prior to 3 uLs of the liquid BEV sample being applied for 30s. Excess sample was wicked away and grids were exposed to three 15 uL washes with molecular grade water prior to negative staining with two applications of 10 μL of filtered 0.75% uranyl formate, with wicking of excess fluid using hardened Whatman 50 filter paper, between steps. The grids were allowed to dry prior to examination on a Talos 120C transmission electron microscope equipped with a CETA 16 megapixel camera (Thermo Fisher) for image capture using TIA (Thermo Fisher).

### Nanoparticle tracking analysis

Nanoparticle size distributions were assessed using nanoparticle tracking analysis, equipped with a sCMOS camera, 532 nm green laser, and a 565 nm long pass filter (NanoSight NS300; Malvern Panalytical, Malvern, United Kingdom). For clinical concentrations, the beta-lactam samples were diluted 1:1000 in PBS. For 2MIC, beta-lactam samples were diluted 1:100 in PBS, and all other samples were diluted 1:10 in PBS. Three measurements of each sample were performed for 30 seconds each, with a camera level of 13 and detection threshold of 5, and average particle concentrations were reported after taking the dilution factors into account.

## Data availability

All data necessary for evaluating the conclusions of this study are present within the manuscript and the supporting information. Additional data (including individual immunoblot images) can be shared upon request.

### Supporting information

This article contains supporting information.

## Acknowledgments

We thank Dr. Kwang Sik Kim (Johns Hopkins Children’s Center) for the gift of *E. col*i K1 RS218. Transmission electron microscopy was conducted by staff in the Electron Microscopy Resource (EMR) in the Center for Advanced Research Technology (CART) at the University of Rochester Medical Center.

## Author contributions

L.V.M., T. G., and P. T. conceptualization; P. T., N. S., J. C., G. G., N. B., M. V., A. M., J. P., and M. D. experimentation; P. T., L. V. M writing, T. G. and A. P. review & editing, L. V. M. and T. G. funding acquisition.

## Funding and additional information

This work was supported by NIAID of the National Institutes of Health under award number R21AI163782 to L. V. M. and T. G. The content is solely the responsibility of the authors and does not necessarily represent the official views of the National Institutes of Health.

## Conflict of interest

The authors declare that they have no conflicts of interest with the contents of this article.

## Abbreviations

The abbreviations used are:

OMV: outer membrane vesicle
BEV: Bacterial extracellular vesicle
PAL: Peptidoglycan associated protein
OmpA: Outer membrane protein
A; NTA: nanoparticle tracking analysis
TEM: transmission electron microscopy

## Supplementary Data

**Figure S1.**
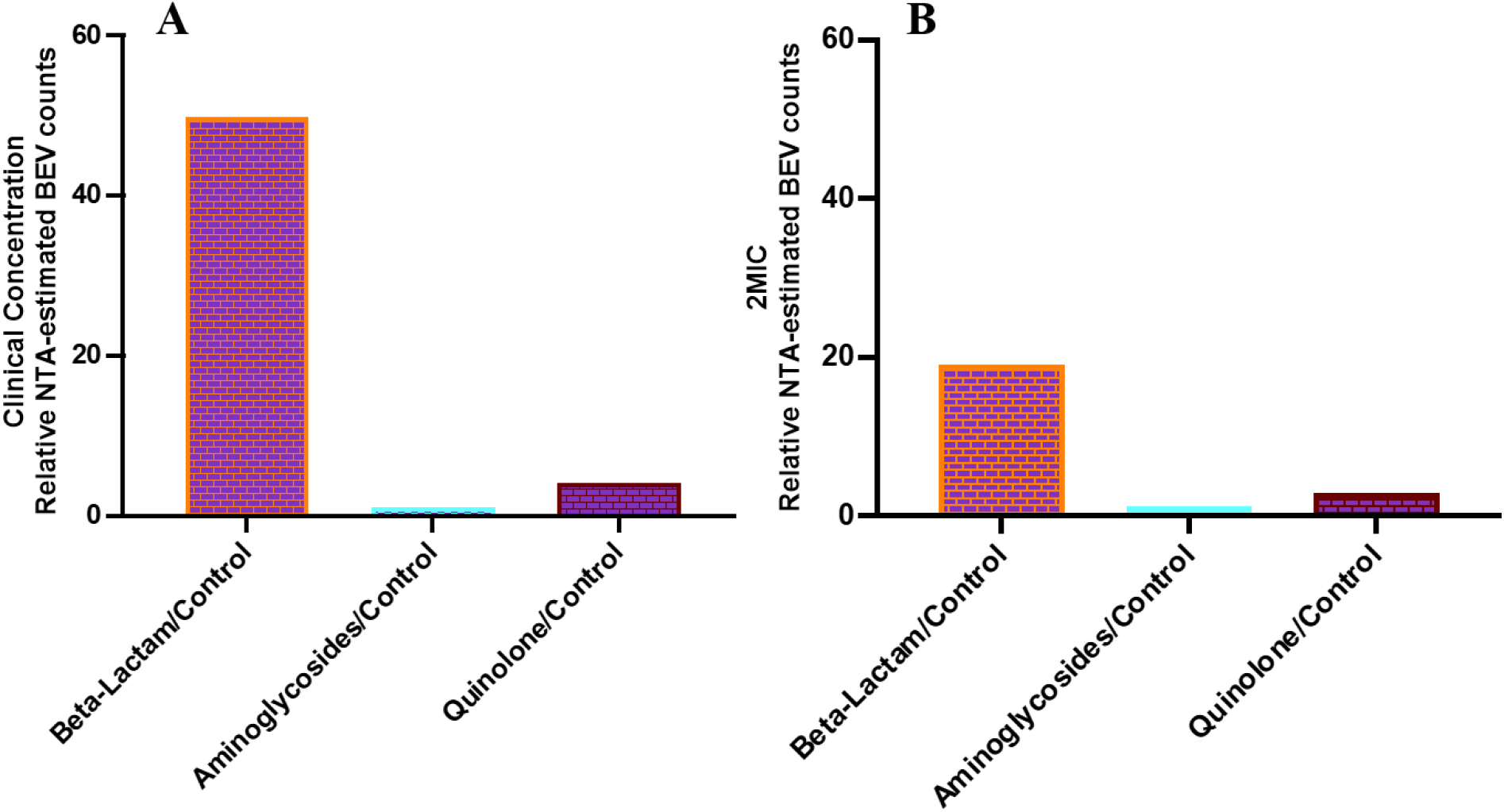
Relative BEV counts determined from average NTA data. Relative BEV counts were determined using the average NTA-derived particle counts for each class of antibiotic compared to control (no antibiotic) for clinical concentrations (A) and 2MIC (B).

**Figure S2.**
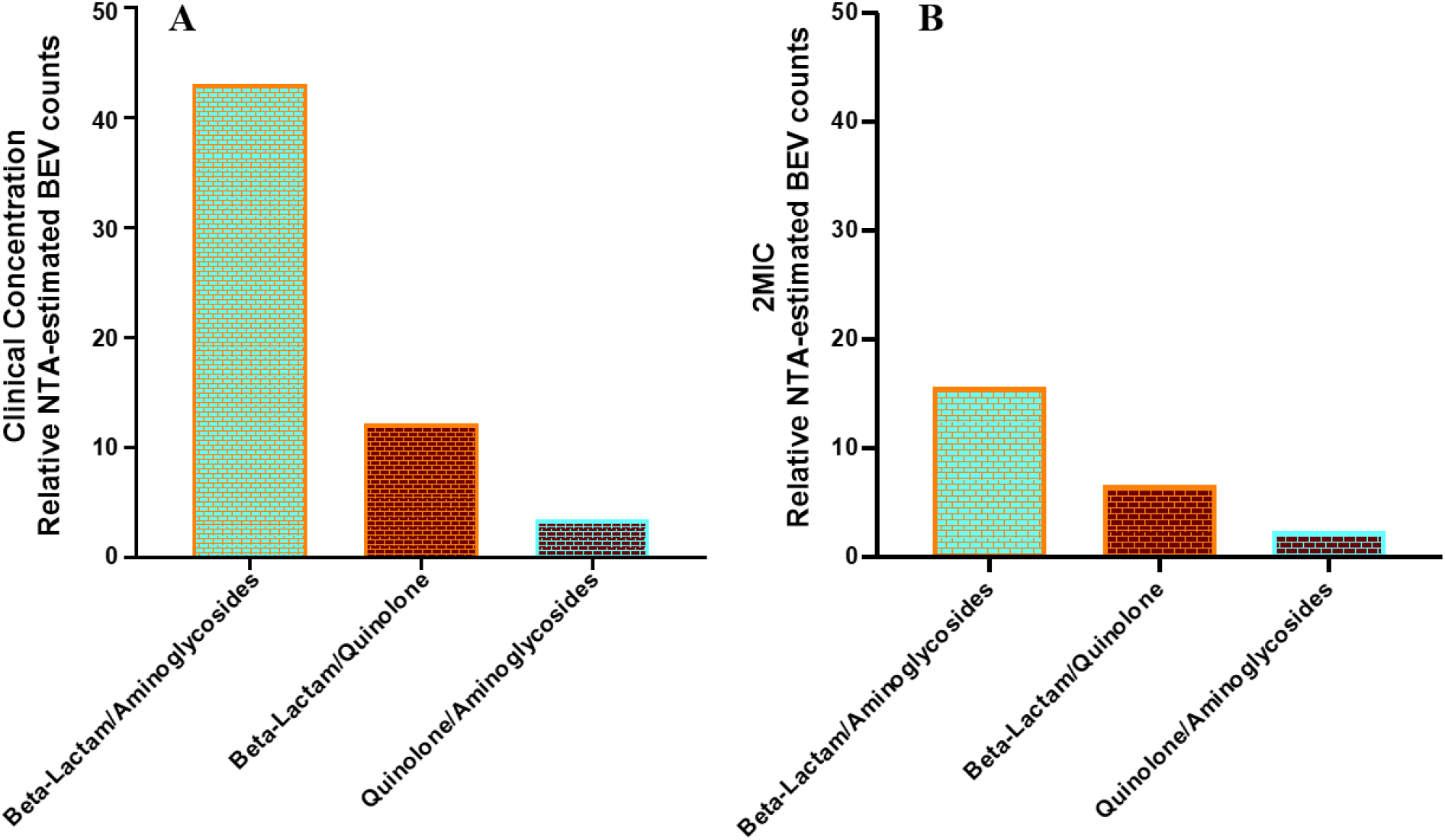
Relative BEV counts determined from average NTA data for beta-lactam antibiotics vs aminoglycosides, beta-lactam antibiotics vs quinolone, and quinolone vs aminoglycosides at clinical concentrations (A) and 2MIC (B).

## Notes

### Competing Interest Statement

The authors have declared no competing interest.

